# Co-expression analysis reveals interpretable gene modules controlled by *trans*-acting genetic variants

**DOI:** 10.1101/2020.04.22.055335

**Authors:** Liis Kolberg, Nurlan Kerimov, Hedi Peterson, Kaur Alasoo

## Abstract

**Background:** Developing novel therapies for complex disease requires better understanding of the causal processes that contribute to disease onset and progression. Although *trans*-acting gene expression quantitative trait loci (*trans*-eQTLs) can be a powerful approach to directly reveal cellular processes modulated by disease variants, detecting *trans*-eQTLs remains challenging due to their small effect sizes and large number of genes tested. However, if a single *trans*-eQTL controls a group of co-regulated genes, then multiple testing burden can be greatly reduced by summarising gene expression at the level of co-expression modules prior to *trans*-eQTL analysis.

**Results:** We analysed gene expression and genotype data from six blood cell types from 226 to 710 individuals. We inferred gene co-expression modules with five methods on the full dataset, as well as in each cell type separately. We detected a number of established co-expression module *trans*-eQTLs, such as the monocyte-specific associations at the *IFNB1* and *LYZ* loci, as well as a platelet-specific *ARHGEF3* locus associated with mean platelet volume. We also discovered a novel *trans* association near the *SLC39A8* gene in LPS-stimulated monocytes. Here, we linked an early-response *cis*-eQTL of the *SLC39A8* gene to a module of co-expressed metallothionein genes upregulated more than 20 hours later and used motif analysis to identify zinc-induced activation of the MTF1 transcription factor as a likely mediator of this effect.

**Conclusions:** Our analysis provides a rare detailed characterisation of a *trans*-eQTL effect cascade from a proximal *cis* effect to the affected signalling pathway, transcription factor, and target genes. This highlights how co-expression analysis combined with functional enrichment analysis can greatly improve the identification and prioritisation of *trans*-eQTLs when applied to emerging cell-type specific datasets.

## Background

Genome-wide association studies have been remarkably successful at identifying genetic variants associated with complex traits and diseases. To enable pharmacological and other interventions on these diseases, linking associated variants to causal intermediate phenotypes and processes is needed. A canonical example is the causal role of circulating LDL cholesterol in cardiovascular disease [1]. However, discovering clinically relevant intermediate phenotypes has so far remained challenging for most complex diseases. At the molecular level, *cis*-acting gene expression quantitative trait loci (*cis*-eQTLs) can be used to identify putative causal genes at disease-associated loci, but due to widespread co-regulation between neighbouring genes [2] and poor understanding of gene function, these approaches often identify multiple candidates whose functional relevance for the disease is unclear.

A promising approach to overcome the limitations of *cis*-eQTLs is *trans*-eQTL analysis linking disease-associated variants via signalling pathways and cellular processes (*trans*-acting factors) to multiple target genes. Although *trans*-eQTLs are widespread [3], most transcriptomic studies in various cell types and tissues are still underpowered to detect them [4]. This is due to limited sample sizes of current eQTL studies, small effect sizes of *trans*-eQTLs, and the large number of tests performed (>10^6^ independent variants with >10^4^ genes). To reduce the number of tested phenotypes, co-expression analysis methods are sometimes used to aggregate individual genes to co-expressed modules capturing signalling pathways and cellular processes [5]. Such approaches have been successful in identifying *trans*-eQTLs in yeast [6] as well as various human tissues [7–9] and purified immune cells [10,11].

Gene co-expression modules can be detected with various methods. Top-down matrix factorisation approaches such as independent component analysis (ICA) [12], sparse decomposition of arrays (SDA) [7] and probabilistic estimation of expression residuals (PEER) [13] seek to identify latent factors that explain large proportion of variance in the dataset. In these models, a single gene can contribute to multiple latent factors with different weights. In contrast, bottom-up gene expression clustering methods such as weighted gene co-expression network analysis (WGCNA) [14] seek to identify non-overlapping groups of genes with highly correlated expression values. Recently, both matrix factorisation and co-expression clustering methods have been further extended to incorporate prior information about biological pathways and gene sets, resulting in pathway-level information extractor (PLIER) [9] and funcExplorer [15], respectively. Out of these methods, ICA, WGCNA, SDA and PLIER have previously been used to find *trans*-eQTLs for modules of co-expressed genes [7–11], but only a single method at a time. However, since different methods solve distinct optimisation problems, they can detect complementary sets of co-expression modules [5]. Thus, applying multiple co-expression methods to the same dataset can aid *trans*-eQTL detection by identifying complementary sets of co-expression modules capturing a wider range of biological processes.

Another aspect that can influence co-expression module detection is how the data is partitioned prior to analysis [5]. This is particularly relevant when data from multiple cell types or conditions is analysed together. When co-expression analysis is performed across multiple cell types or conditions, then the majority of detected gene co-expression modules are guided by differential expression between cell types [16,17]. As a result, cell-type-specific co-expression modules can be missed due to weak correlation in other cell types [16]. One strategy to recover such modules is to perform co-expression analysis in each cell type separately [5].

In this study, we performed comprehensive gene module *trans*-eQTL analysis across six major blood cell types and three stimulated conditions from five published datasets. To maximise gene module detection, we applied five distinct co-expression analysis methods (ICA, PEER, PLIER, WGCNA, funcExplorer) to the full dataset as well as individual cell types and conditions separately. Using a novel aggregation approach based on statistical fine mapping, we grouped individual *trans*-eQTLs to a set of non-overlapping loci. Extensive follow-up with gene set and transcription factor motif enrichment analyses allowed us to gain additional insight into the functional impact of *trans*-eQTLs and prioritise loci for further analyses. In addition to replicating two known monocyte-specific *trans*-eQTLs at the *IFNB1 [11,17–19]* and *LYZ* loci [10,20,21], we found that the *trans*-eQTL at the *ARHGEF3* locus detected in multiple whole blood datasets [8–10,22] was highly specific to platelets in our analysis. Finally, we also detected a novel association at the *SLC39A8* locus that controlled a group of genes encoding zinc-binding proteins in LPS-stimulated monocytes.

## Results

### Cell types, conditions and samples

We used gene expression and genotype data from five previously published studies from three independent cohorts [18,20,23–25]. The data consisted of CD4+ and CD8+ T cells [23,24], B cells [20,23], neutrophils [23,25], platelets [23], naive monoctyes [18,23] and monocytes stimulated with lipopolysaccharide for 2 or 24 hours (LPS 2h, LPS 24h) and interferon-gamma for 24 hours (IFNγ 24h) [18]. The sample size varied from n = 226 in platelets to n = 710 in naive monocytes (Figure 1A). After quality control, normalisation and batch correction (see “Methods”), the final dataset consisted of 18,383 unique protein coding genes profiled in 3,938 samples from 1,037 unique genotyped individuals of European ancestries (Figure 1B). Even though the samples originated from five different studies, they clustered predominantly by cell type of origin (Figure 1B).

**Figure 1.**
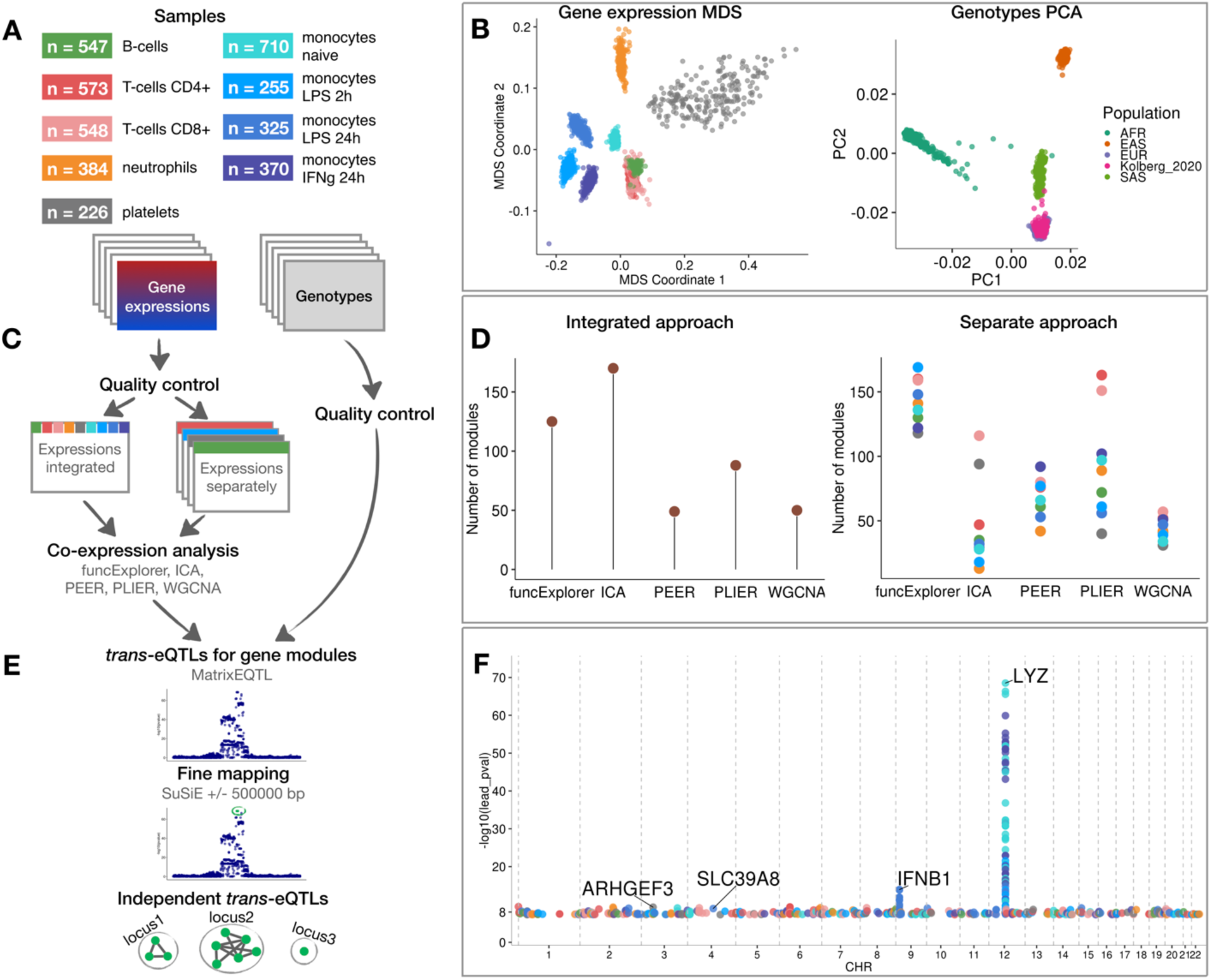
Data, analysis workflow and results. **A** Sample sizes of cell types and conditions included in the analysis. LPS - lipopolysaccharide, IFNg - interferon-gamma. **B** Multidimensional scaling (MDS) analysis of the gene expression data and principal component analysis (PCA) of genotype data after quality control and normalisation. Cell types and conditions are color-coded according to panel A. Genotyped samples from this study have been projected to the 1000 Genomes Project reference populations. **C** Following quality control, five co-expression methods were applied to two different data partitioning approaches: (1) gene expression profiles across all cell types and conditions were analysed together (integrated approach), (2) gene expression profiles from each cell type and condition were analysed separately (separate approach). **D** The number of gene modules detected from integrated and separate analyses. **E** For *trans*-eQTL analysis we used the estimated module activity profile (‘eigengene’) as our phenotype. To identify independent *trans*-eQTLs, we performed statistical fine mapping for all nominally significant (P-value < 5×10^-8^) associations and grouped together all associations with overlapping credible sets. **F** Manhattan plot of nominally significant (P-value < 5×10^-8^) *trans*-eQTLs. Each point corresponds to a gene module that was associated with the corresponding locus and is color-coded by the cell type from panel A.

### Detecting *trans*-eQTLs regulating modules of co-expressed genes

We performed co-expression analyses with ICA, WGCNA, PLIER, PEER and funcExplorer on the full gene expression dataset (integrated approach) as well as on each cell type and condition separately (separate approach) (Figure 1C). In total, we obtained 482 gene modules from the integrated approach and 3,509 from the separate clustering of different cell types (Figure 1D; Additional file 1: Figure S1). For every module, the methods inferred a single characteristic expression pattern (‘eigengene’) that represents the expression profiles of the module genes across the samples. Although implementation details varied between methods (see “Methods”), these eigengene profiles were essentially linear combinations of expression levels of genes belonging to the modules.

For *trans*-eQTL analysis, we included 6,861,056 common (minor allele frequency > 5%) genetic variants passing strict quality control criteria. First, we used linear regression implemented in MatrixEQTL [26] package to identify all genetic variants nominally associated (P-value < 5×10^-8^) with the eigengenes of each of the 3,991 co-expression modules detected across 9 cell types and conditions. We performed *trans*-eQTL analysis in each cell type and condition separately. Next, we used SuSiE [27] to fine map all significant associations to 864 independent credible sets of candidate causal variants (Figure 1E). Since we applied five co-expression methods to both integrated and cell type specific (separated) datasets, we found a large number of overlapping genetic associations. We thus aggregated overlapping credible sets from 864 associations to 601 non-overlapping genomic loci (Additional file 1: Figure S2; see “Methods”). We observed that some, especially smaller, co-expression modules were driven by strong *cis*-eQTL effects that were controlling multiple neighbouring genes in the same module. To exclude such effects, we performed gene-level eQTL analysis for 18,383 protein-coding genes and the 601 lead variants identified above. We excluded co-expression modules where the module lead variant was not individually associated with any of the module genes in *trans* (> 5Mb away) (see “Methods”). This step reduced the number of nominally significant *trans*-eQTL loci to 303 (Figure 1F; Additional file 2). Finally, to account for the number of co-expression modules tested, we used both Benjamini-Hochberg false discovery rate (FDR) and Bonferroni correction (see “Methods”). The FDR 10% threshold reduced the number of significant associations to 140 and Bonferroni threshold retained only 4 significant loci, including loci near *IFNB1* (Additional file 1: Figure S3) and *LYZ* (Additional file 1: Figure S4) genes that have been previously reported in several other studies [10,17–21] (Additional file 1: Table S1). While the strong *trans*-eQTL signals at the *IFNB1* and *LYZ* loci were detected by all co-expression methods in both integrated and separate analyses, most associations were detected by only a subset of the analytical approaches (Additional file 2). Furthermore, almost all *trans*-eQTLs that we detected had highly cell type and condition specific effects (Additional file 1: Figure S5). We will now dissect two such cell type and condition specific associations in more detail.

### Platelet specific *trans*-eQTL at the *ARHGEF3* locus is associated with multiple platelet traits

We found that the rs1354034 (T/C) variant located within the *ARHGEF3* gene is associated with three co-expression modules in platelets: one ICA module detected in integrated analysis (IC68, 1,074 genes) and two co-expression modules detected in a platelet-specific analysis by PLIER (X6.WIERENGA_STAT5A_TARGETS_DN, 918 genes) and funcExplorer (Cluster_12953, 5 genes) (Figure 2B, Additional file 1: Figure S6). The T allele increases the expression of the *ARHGEF3* gene in *cis* and the two lead variants are the same (Figure 2A). Furthermore, both the *cis* and *trans*-eQTLs colocalise with a GWAS hit for mean platelet volume (*cis* PP4 = 0.99, *trans* PP4 > 0.99 for all modules), platelet count (*cis* PP4 = 0.99, *trans* PP4 > 0.99 for all modules) and plateletcrit (*trans* PP4 > 0.99 for all modules) (Figure 2A) [28]. Interestingly, *ARHGEF3* itself is not in any of the three modules and the module eigengenes are not strongly co-expressed with *ARHGEF3* (Pearson’s r ranging from 0.07 to 0.33 in platelets). While IC68 and X6.WIERENGA_STAT5A_TARGETS_DN share 74 overlapping genes (one-sided Fisher’s exact test P-value = 0.003), none of the genes in Cluster_12953 is in any of the other modules.

Although the *ARHGEF3 trans*-eQTL has been detected in multiple whole blood *trans*-eQTL studies [3,8,9,22] (Additional file 1: Table S1), our analysis demonstrates that this association is highly specific to platelets and not detected in other major blood cell types (Figure 2B). Furthermore, even though *ARHGEF3* is expressed in multiple cell types, the *cis*-eQTL effect is also only visible in platelets (Additional file 1: Figure S6). Reassuringly, the *trans*-eQTL effect sizes in our small platelet sample (n = 216) are correlated (Pearson’s r = 0.68, P-value = 5.1×10^-12^) with the effects from the largest whole blood *trans*-eQTL meta-analysis [3] (n = 31,684) (Additional file 1: Figure S7). The platelet specificity of the *ARHGEF3* association is further supported by functional enrichment analysis with g:Profiler [29] which found that both the PLIER module X6.WIERENGA_STAT5A_TARGETS_DN and target genes from the gene-level analysis were strongly enriched for multiple terms related to platelet activation (Figure 2E; https://biit.cs.ut.ee/gplink/l/rtyNe5M2R4). Cluster_12953, however, was enriched for cellular response to iron ion, suggesting that *ARHGEF3* might be involved in multiple independent processes [9,30]. Altogether, these results demonstrate how a *trans*-eQTL detected in whole blood can be driven by a strong signal present in only one cell type.

**Figure 2.**
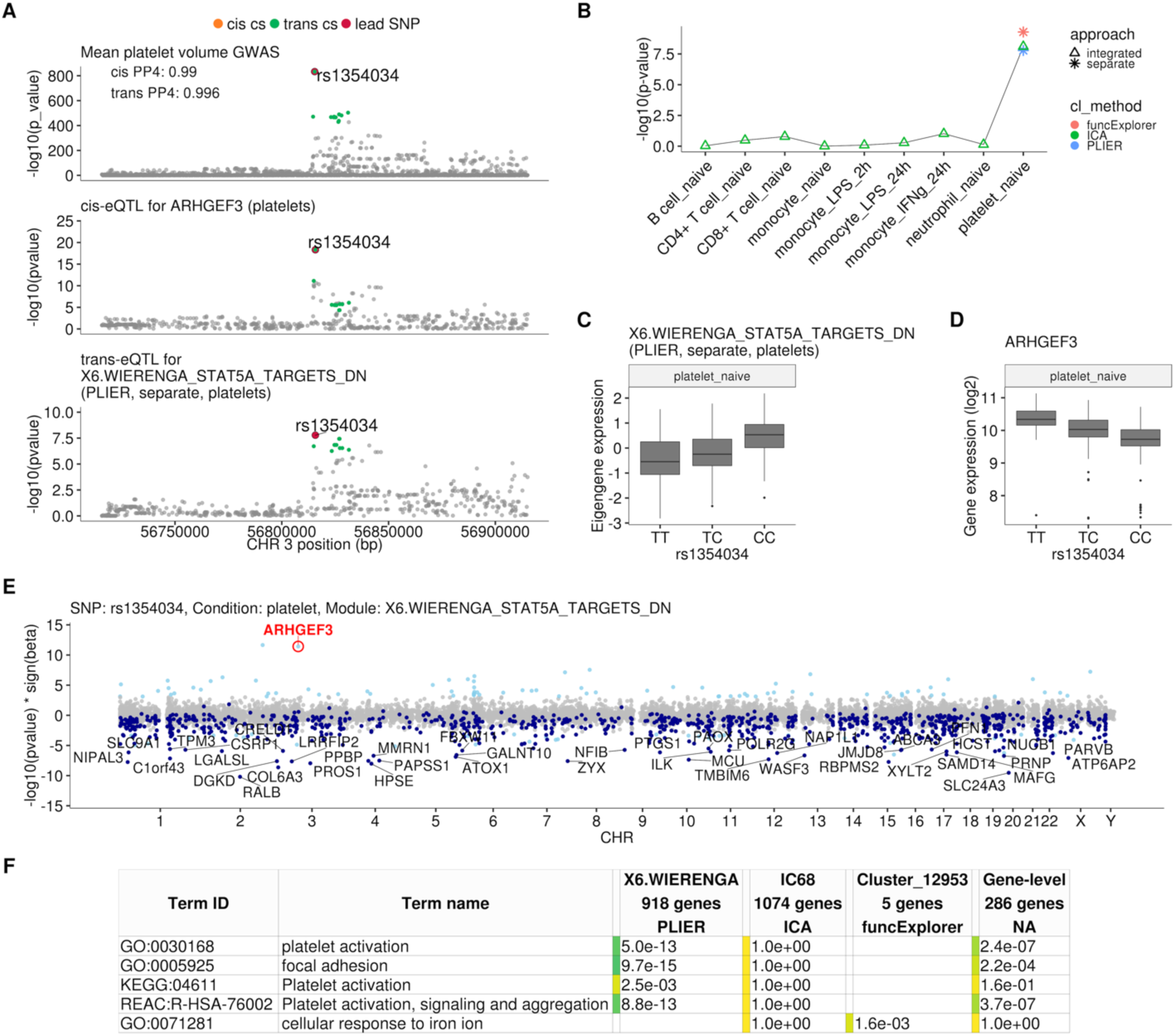
Platelet-specific *trans*-eQTL at the *ARHGEF3 locus*. **A** Regional plots showing colocalisation between GWAS signal for mean platelet volume [28], *cis*-eQTL for *ARHGEF3* in platelets and *trans*-eQTL for a platelet-specific co-expression module detected by PLIER. *Cis* and *trans* credible sets (cs) are marked on the plots. The *cis* credible set consists of only the lead variant. **B** Line graph showing that the association between the modules and *ARHGEF3* locus is platelet specific. In cell-type-specific clustering, only a single P-value from the corresponding cell type is available. The integrated modules have P-values from each of the cell types and the values are connected by a line. **C** Association between the *trans*-eQTL lead variant (rs1354034) and eigengene of module X6.WIERENGA_STAT5A_TARGETS_DN in platelets. **D** Association between the *trans*-eQTL lead variant (rs1354034) and *ARHGEF3* expression in platelets. **E** Manhattan plot of gene-level eQTL analysis for the *trans*-eQTL lead variant. Dark blue points highlight the genes in module X6.WIERENGA_STAT5A_TARGETS_DN. **F** Functional enrichment analysis of modules associated with *ARHGEF3* locus (see full results at https://biit.cs.ut.ee/gplink/l/rtyNe5M2R4). Empty cell indicates that no gene in the module is annotated to the corresponding term, enrichment P-value = 1 shows that at least some of the genes in the module are annotated to the term, but not enough to report over-representation. The last column combines the FDR 5% significant genes from the gene-level analysis. GO - Gene Ontology, KEGG - Kyoto Encyclopedia of Genes and Genomes Pathways, REAC - Reactome Pathways.

### *SLC39A8* locus is associated with zinc ion homeostasis in LPS-stimulated monocytes

One of the novel results in our analysis was a locus near the *SLC39A8* gene that was associated (P-value = 1.2×10^-9^) with a single co-expression module detected by funcExplorer (Cluster_10413) in monocytes stimulated with LPS for 24 hours (Figure 3A-C). The module consisted of five metallothionein genes (*MT1A, MT1F, MT1G, MT1H, MT1M*) all located in the same locus on chromosome 16 (Figure 3D). Although the *trans*-eQTL lead variant (rs75562818) was significantly associated with the expression of the *SLC39A8* gene (Figures 3A and 3D), the two association signals did not colocalise and the credible sets did not overlap (Figure 3A; Additional file 1: Figure S8), indicating that the *cis*-eQTL detected in naive and stimulated monocytes in our dataset is not the main effect driving the *trans*-eQTL signal. Furthermore, the expression of *SLC39A8* was only moderately correlated with the eigengene value of Cluster_10413 (Pearson’s r = 0.27). Since *SLC39A8* is strongly upregulated (log2 fold-change = 3.53) in response to LPS already at two hours (Figure 4A), we speculated that there might be a transient eQTL earlier in the LPS response. To test this, we downloaded the eQTL summary statistics from the Kim-Hellmuth *et al*, 2017 study that had mapped eQTLs in monocytes stimulated with LPS for 90 minutes and six hours [31]. Indeed, we found that the *cis*-eQTL 90 minutes after LPS stimulation colocalised with our *trans*-eQTL (Figure 3A) and this signal disappeared by six hours after stimulation (Additional file 1: Figure S9).

To understand the function of the *SLC39A8* locus, we turned to the target genes. Gene-level analysis identified two more metallothionein genes (*MT1E* and *MT1X*) from the same locus as likely target genes (Figure 3D). Enrichment analysis with g:Profiler revealed that these genes were enriched for multiple Gene Ontology terms and pathways related to zinc ion homeostasis (Figure 3E, full results at https://biit.cs.ut.ee/gplink/l/v7FhJRn4Rj). Furthermore, the promoter regions of the 7 genes were also enriched for the binding motif of the metal transcription factor 1 (MTF1) transcription factor (P-value = 2.1×10^-4^, Figure 3E). Taken together, these results suggest that a transient eQTL of the *SLC39A8* gene 90 minutes after stimulation regulates the expression of 7 zinc binding proteins 24 hours later. Multiple lines of literature evidence support this model (Figure 4B). First, the ZIP8 protein coded by the *SLC39A8* gene is a manganese and zinc ion influx transporter [32]. Secondly, *SLC39A8* is upregulated by the NF-κB transcription factor in macrophages and monocytes in response to LPS and this upregulation leads to increased intracellular Zn^2+^ concentration [33]. Third, Zn^2+^ influx increases the transcriptional activity of the metal transcription factor 1 (MTF1) [34] and metallothioneins, which act as Zn^2+^-storage proteins, are well known target genes of the MTF1 transcription factor [35]. Finally, *SLC39A8* knockdown in mice leads to decreased expression of the metallothionein 1 (*MT1*) gene [33].

**Figure 3.**
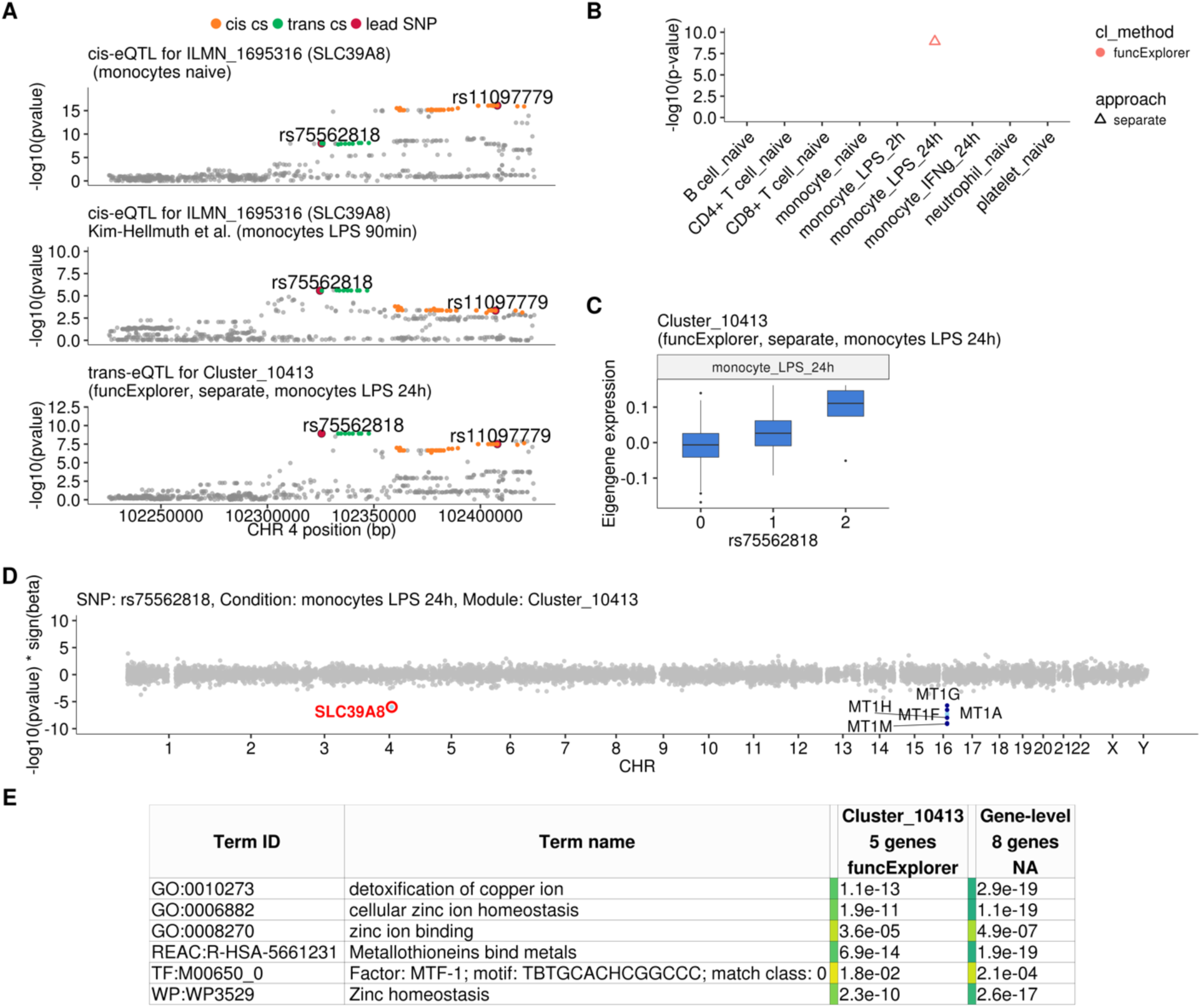
Transient *cis*-eQTLs for *SLC39A8* is associated with the expression of seven metallothionein genes in *trans* in monocytes stimulated with LPS for 24 hours. **A** Regional plots comparing association signals between naive (rs11097779) and transiently induced *cis*-eQTLs (rs75562818) for *SLC39A8* and *trans*-eQTL (rs75562818) for a module of five co-expressed metallothionein genes. LPS-induced *cis*-QTL summary statistics 90 minutes post stimulation (n = 134) were obtained from Kim-Hellmuth et al, 2017 [31]. **B** Graph showing that the association between the module and *SLC39A8* locus is stimulation specific. As this module was detected by a cell type specific clustering, only a single value from the corresponding cell type is available. **C** Association between *trans*-eQTL (rs75562818) and eigengene of funcExplorer module Cluster_10413 in monocytes after 24 hours of LPS stimulation. **D** Manhattan plot of gene-level eQTL analysis for rs75562818. Dark blue points highlight the genes in module Cluster_10413. **E** Functional enrichment analysis of the *SLC39A8* associated module (see https://biit.cs.ut.ee/gplink/l/v7FhJRn4Rj for full results). The last column combines the FDR 5% significant genes from the gene-level analysis. MTF1 - metal transcription factor 1. GO - Gene Ontology, WP - WikiPathways, REAC - Reactome Pathways, TF - transcription factor binding sites from TRANSFAC.

To see if the *SLC39A8 trans*-eQTL might be associated with any higher level phenotypes, we queried the GWAS Catalog [36] database with the ten variants from the *trans*-eQTL 95% credible set. We found that a lead variant for red blood cell distribution width (rs7692921) was one of the variants in our credible set and in high LD (r^2^ = 0.991) with the *trans*-eQTL lead variant (Figure 4C) [37]. However, neither of the eQTL variants was in LD with a known missense variant (rs13107325) in the *SLC39A8* gene that has been associated with schizophrenia, Parkinson’s disease and other traits (Figure 4C) [38].

**Figure 4.**
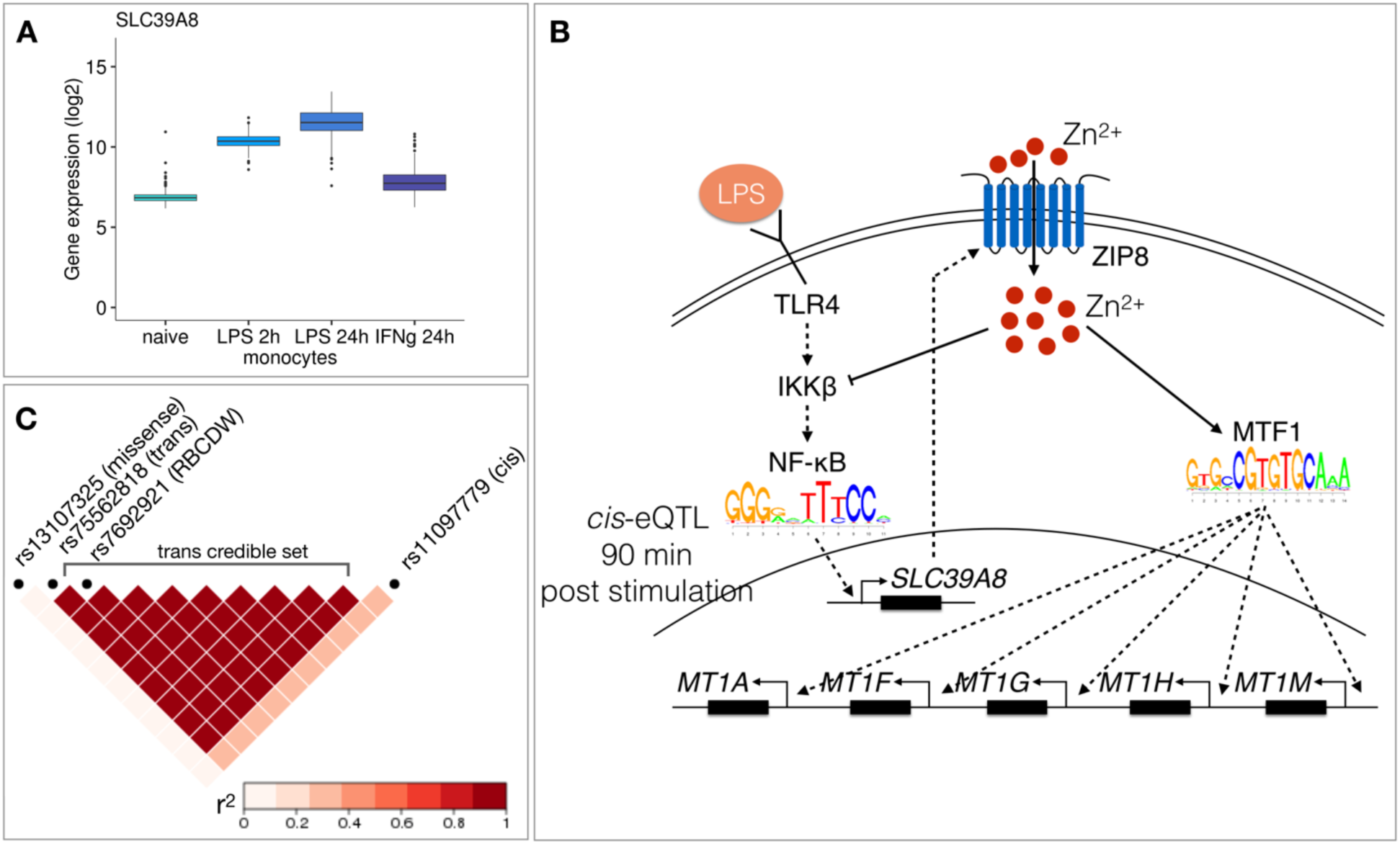
Molecular mechanisms underlying the *SLC39A8 trans*-eQTL locus. **A** *SLC39A8* gene expression values (log_2_ intensities) across naive and stimulated monocytes. **B** Overview of the known regulatory interactions underlying the *cis* and *trans* eQTL effects at the *SLC39A8* locus. Figure adapted from [33]. **C** Pairwise LD (r^2^ within 1000 Genomes European populations) between the *SLC39A8* variants highlighting missense variant (rs13107325), *trans*-eQTL (rs75562818), red blood cell distribution width (RBCDW) associated SNP (rs7692921) in our credible set and the *cis* lead variant from naive monocytes (rs11097779). LD was calculated using the LDlinkR (v.1.0.2) R package [39].

## Discussion

Given that *trans*-eQTLs have been more difficult to replicate between studies and false positive associations can easily occur due to technical issues [40,41], it is increasingly important to effectively summarise and prioritise associations for follow up analyses and experiments. We found that aggregation of credible sets of eigengene profiles from multiple co-expression methods (Additional file 1: Figure S2) successfully reduced the number of independent associations, but this still retained more than 300 loci that we needed to evaluate. To further prioritise associations, we used gene set and transcription factor motif enrichment analysis of the *trans*-eQTL target genes. Although motif analysis is often underpowered, it can provide directly testable hypotheses about the *trans*-eQTL mechanism such as the MTF1 transcription factor that we identified at the *SLC39A8* locus. Similar approaches have also been successfully used to characterise *trans*-eQTLs involving IRF1 and IRF2 transcription factors [18,42].

A major limitation of co-expression based approach for *trans*-eQTL mapping is that many true co-expression modules can remain undetected by various co-expression analysis methods. We sought to overcome this by aggregating results across five complementary co-expression methods. We found that while all methods were able to discover strong co-expression module *trans*-eQTLs such as those underlying the *IFNB1* (Additional file 1: Figure S3) and *LYZ* (Additional file 1: Figure S4) associations, most co-expression module *trans*-eQTLs were only detected by a subset of the analysis methods. For example, the *ARHGEF3* association was detected by three of the five methods (Figure 2B) and *SLC39A8* co-expression module was found only by funcExplorer and only when samples from LPS-stimulated monocytes were analysed separately (Figure 3B). While it is always possible to include additional analysis methods, this should be appropriately weighed against the increase in the number of phenotypes tested. In our analysis, we decided to first use a relaxed nominal significance threshold of P-value < 5×10^-8^ and subsequently focus on associations that we could either replicate in independent datasets or find significant support from the literature. Finally, if the *trans*-eQTL locus controls a single or a small number of genes then co-expression-based approaches are probably not well suited to detect such associations and gene-level analysis is still required.

Since eQTL datasets from purified cell types are still relatively small and single cell eQTL datasets are even smaller [43], it is tempting to perform *trans*-eQTL analysis on whole tissue datasets such as the brain or whole blood [3]. However, it remains unclear what fraction of cell type and condition specific *trans*-eQTLs can be detected in whole tissue datasets collected from healthy donors. Although we were able to replicate the *ARHGEF3* association in the eQTLGen whole blood meta-analysis, because our fine mapped lead variant happened to be one of the 10,317 variants tested in eQTLGen, systematic replication requires genome-wide summary statistics that are currently lacking for *trans*-eQTL analyses. Secondly, tissue datasets can be biased by cell type composition effects. These can lead to spurious *trans*-eQTL signals, because genetic variants associated with cell type composition changes would appear as *trans*-eQTLs for cell-type-specific genes [3]. Furthermore, multiple studies have demonstrated that also the co-expression signals in tissues are largely driven by cell type composition effects [44–46]. Thus, even though PLIER detected the ARHGEF3 *trans*-eQTL in whole blood, this could have been at least partially driven by the change in platelet proportion between individuals [9]. Our analysis in purified cell types enabled us to verify that this was a truly platelet-specific genetic association.

Although both in the case of *ARHGEF3* and *SLC39A8,* we could be reasonably confident that the expression level of the *cis* gene mediated the observed *trans*-eQTL effect, there was only a modest correlation (Pearson’s r between 0.07 and 0.33) between the *cis* gene expression and the corresponding *trans* co-expression module expression. In case of *SLC39A8* there seemed to be a temporal delay with the *cis*-eQTL being active early in LPS response and *trans*-eQTL appearing much later after proposed accumulation of the ZIP8 protein and increase in intracellular zinc concentration. Temporal delay has also been reported for the *trans*-eQTLs at the *INFB1* [18] and *IRF1* [42] loci. Similarly, stimulation-specific *cis*-eQTL at *CR1* locus in whole blood is not able to explain the full extent of the *trans*-eQTL observed at the same locus [47]. This suggests that if *cis* and *trans* effects are separated from each other either in time (early *versus* late response) or space (different cell types that interact with each other), then this might limit the power of methods that rely on genetically predicted gene expression levels to identify regulatory interactions [22,48,49] and infer causal models. This can also have a negative impact on mediation analysis [50–52], which seeks to estimate the proportion of *trans*-eQTL variance explained by the expression level of the *cis* gene. Altogether, these results indicate that limiting *trans*-eQTL analysis to missense variants and to variants that have been detected as *cis*-eQTLs in the same cell type might miss some true associations, because the *cis* effect might be active in some other, yet unprofiled, context.

## Conclusions

We have performed a large-scale *trans*-eQTL analysis in six blood cell types and three stimulated conditions. We demonstrate that co-expression module detection combined with gene set enrichment analysis can help to identify interpretable *trans*-eQTLs, but these results depend on which co-expression method is chosen for analysis and how the input data is partitioned beforehand. We find that the detected *trans*-eQTLs are highly cell type specific and we use these approaches to perform in-depth characterisation of two cell type specific *trans*-eQTL loci: platelet-specific *trans*-eQTL near the *ARHGEF3* gene and monocyte-specific associations near the *SLC39A8* locus. In both cases, the co-expression modules were enriched for clearly interpretable Gene Ontology terms and pathways, which directly guided literature review and more detailed analyses. We believe that applying co-expression and gene set enrichment based approaches to larger eQTL datasets has the power to detect many more additional associations while simultaneously helping to prioritise *trans*-eQTLs for detailed experimental or computational characterisation.

## Methods

### Datasets used in the analysis

#### CEDAR

The CEDAR dataset [23] contained gene expression and genotype data from CD4+ T-cells, CD8+ T-cells, CD19+ B-cells and CD14+ monocytes, CD15+ neutrophils and platelets from up to 323 individuals. The raw gene expression data generated with Illumina HumanHT-12 v4 arrays were downloaded from ArrayExpress [53] (accession E-MTAB-6667). The raw IDAT files were imported into R using the `readIdatFiles` function from the beadarray v2.28 [54] Bioconductor package.

The raw genotype data generated by Illumina HumanOmniExpress-12 v1_A genotyping arrays were also downloaded from ArrayExpress (accession E-MTAB-6666). Genotype calling was performed with Illumina GenomeStudio v2.0.4, after which the raw genotypes were exported in PLINK format.

#### Kasela, 2017

Kasela *et al*, 2017 [24] generated gene expression and genotype data from CD4+ and CD8+ T cells from 297 unique donors. The raw gene expression data generated with Illumina HumanHT-12 v4 arrays were downloaded from Gene Expression Omnibus (accession GSE78840). The genotype data generated by Illumina HumanOmniExpress-12 v1_A genotyping arrays were obtained from the Estonian Genome Center, University of Tartu (https://www.geenivaramu.ee/en/biobank.ee/data-access). Ethical approval was obtained from the Research Ethics Committee of the University of Tartu (approval 287/T-14).

#### Fairfax, 2012, Fairfax, 2014 and Naranbhai, 2015

Fairfax et al, 2012 [20] profiled gene expression in CD19+ B cells from 282 individuals (ArrayExpress accession E-MTAB-945). Fairfax et al, 2014 [18] profiled gene expression in naive CD14+ monocytes as well as in cells stimulated with lipopolysaccharide (LPS) for 2 or 24 hours and interferon-gamma for 24 hours from up to 414 individuals (accession E-MTAB-2232). Naranbhai *et al.*, 2015 [25] profiled gene expression in CD15+ neutrophils from 93 individuals (accession E-MTAB-3536). The genotype data for all three studies were generated by Illumina HumanOmniExpress-12 genotyping arrays and were downloaded from European Genome-phenome Archive (accessions EGAD00010000144 and EGAD00010000520).

### Genotype data quality control and imputation

We started with raw genotype data from each study in PLINK format and GRCh37 coordinates. Before imputation, we performed quality control independently on each of the three datasets. Briefly, we used Genotype harmonizer [55] v1.4.20 to align the alleles with the 1000 Genomes Phase 3 reference panel and exclude variants that could not be aligned. We used PLINK v1.9.0 to convert the genotypes to VCF format and used the fixref plugin of the bcftools v1.9 to correct any strand swaps. We used ‘bcftools norm --check-ref x’ to remove any remaining variants where the reference allele did not match the GRCh37 reference genome. Finally, we excluded variants with Hardy-Weinberg equilibrium P-value > 10^-6^, missingness > 0.05 and MAF < 0.01. We also excluded samples with more than 95% of the variants missing. Finally, we merged genotype data from all three studies into a single VCF file.

After quality control, we included 580,802 autosomal genetic variants from 1,041 individuals for imputation. We used a local installation of the Michigan Imputation Server v1.2.1 [56] to perform phasing and imputation with EAGLE v2.4 [57] and Minimac4 [56]. After imputation, we used CrossMap.py v2.8.0 [58] to convert genotype coordinates to GRCh38 reference genome. We used bcftools v1.9.0 to exclude genetic variants with imputation quality score R^2^ < 0.4 and minor allele frequency (MAF) < 0.05. We used PLINK [59] v1.9.0 to perform LD pruning of the genetic variants and LDAK [60] to project new samples to the principal components of the 1000 Genomes Phase 3 reference panel [61]. The Nextflow pipelines for genotype processing and quality control are available from GitHub (https://github.com/eQTL-Catalogue/genotype_qc).

### Detecting sample swaps between genotype and gene expression data

We used Genotype harmonizer [55] v1.4.20 to convert the imputed genotypes into TRITYPER format. We used MixupMapper [62] v1.4.7 to detect sample swaps between gene expression and genotype data. We detected 155 sample swaps in the CEDAR dataset, most of which affected the neutrophil samples. We also detected one sample swap in the Naranbhai, 2015 dataset.

### Gene expression data quality control and normalisation

As a first step, we performed multidimensional scaling (MDS) and principal component analysis (PCA) on each dataset separately to detect and exclude any outlier samples. This was done after excluding the replicate samples and the samples that did not pass the genotype data quality control. Additional outliers were detected after quantile normalisation and adjusting for batch effects. The normalisation was performed using the `lumiN` function from the lumi v.2.30.0 R package [63]. Batch effects, where applicable, were adjusted for with the `removeBatchEffect` function from the limma v.3.34.9 R package [64]. After quality control to exclude outlier samples, the quantile normalised log_2_ intensity values from all datasets were combined. This was followed by regressing out dataset specific batch effects. Only the intensities of 30,353 protein-coding probes were used. Finally, the probe sets were mapped to genes. For genes with more than one corresponding probe set, the probe with the highest average expression was used. 18,383 protein-coding genes with unique Ensembl identifiers remained for co-expression analysis. We did not regress out any principal components from the gene expression data, as this can introduce false positives in *trans*-eQTL analysis due to collider bias [41]. In total, 3,938 samples remained after the quality control (Table 1).

**Table 1.** Number of samples included in the analysis from each study and each cell type.

| Cell type | Fairfax_2012 | Fairfax_2014 | Naranbhai_2015 | Kasela_2017 | CEDAR |
| --- | --- | --- | --- | --- | --- |
| B cell | 281 | - | - | - | 266 |
| T cell CD4+ | - | - | - | 279 | 294 |
| T cell CD8+ | - | - | - | 267 | 281 |
| neutrophil | - | - | 93 | - | 291 |
| platelet | - | - | - | - | 226 |
| monocyte naive | - | 420 | - | - | 290 |
| monocyte LPS 2h | - | 255 | - | - | - |
| monocyte LPS 24h | - | 325 | - | - | - |
| monocyte IFNγ 24h | - | 370 | - | - | - |

### Co-expression analysis

We applied five different methods to identify modules of co-expressed genes from the gene expression data. We used an expression matrix where rows correspond to genes and columns to individuals/samples as input for the methods. The gene expression profiles were centred and standardised prior to analysis. All methods infer gene co-expression modules, each of which can be described by a single expression profile (‘eigengene’) that captures the collective behavior of corresponding genes in the module. The approaches for defining eigengenes differ across the methods (see below). These eigengenes are treated as quantitative traits in the *trans*-eQTL analysis. To detect potential cell type and condition specific modules, we applied the same methods also to the expression matrices from each of the nine cell types and conditions separately. Summaries of the co-expression analysis results from both integrated and cell-type-specific expression data are shown in Additional file 1: Figure S1.

#### Co-expression clustering methods

##### Weighted gene co-expression network analysis (WGCNA)

The WGCNA method [14] identifies non overlapping co-expressed gene modules. Each of the modules is represented by its first principal component of expression values of genes in the module termed as module eigengene. We used the function `blockwiseModules` for automatic block-wise network construction and module identification with default parameters from the dedicated R package WGCNA (v.1.66). The number of modules was detected automatically by the algorithm.

##### funcExplorer

FuncExplorer [15] is a web tool that performs hierarchical clustering on gene expression values which is followed by automated functional enrichment analysis to derive the most biologically meaningful gene modules from the dendrogram. The expression data were uploaded to funcExplorer and the modules were detected using the following parameters: best annotation strategy, P-value threshold 0.01 for enrichment of Gene Ontology, KEGG and Reactome annotations. Every funcExplorer gene module is characterised by the eigengene profile which, like in WGCNA, is the first principal component of module expression values calculated in the same way as in WGCNA. The number of modules is detected automatically by funcExplorer and the different modules consist of non-overlapping sets of genes. The co-expression analysis results are available for browsing from https://biit.cs.ut.ee/funcexplorer/user/2a29dfa6de6b8b733f665352735adaf5 where the option ‘Dataset’ includes the full selection of expression data used in this analysis. Dataset ‘Merged_ENSG_expression’ incorporates integrated samples from all cell types and conditions, ‘CL_0000233_naive’ stands for platelets, ‘CL_0000236_naive’ for B-cells, ‘CL_0000624_naive’ for CD4+ T-cells, ‘CL_0000625_naive’ for CD8+ T-cells, ‘CL_0000775_naive’ for neutrophils, ‘CL_0002057_naive’ for monocytes and ‘CL_0002057_IFNg_24h’, ‘CL_0002057_LPS_24h’, ‘CL_0002057_LPS_2h’ include gene expression matrices from corresponding stimulated monocyte samples.

#### Matrix factorisation methods

Matrix factorisation methods, such as ICA, PLIER and PEER, deconvolve the input gene expression matrix into two related matrices [5]. One of the matrices is the matrix of factor loadings for each sample and the other describes the gene-level weights of the factors. In the case of ICA, PLIER and PEER, we used the factor loadings as module eigengene profiles. For g:Profiler enrichment analysis we used the gene-level weights to define the genes that characterise the modules by choosing the ones that are the most influenced, i.e. the genes at both extremes of gene weight values (2 standard deviations from the mean weights in this module). Thus, different modules can include overlapping sets of genes.

##### Independent component analysis (ICA)

The ICA [12] method attempts to decompose gene expression measurements into independent components (factors) which represent underlying biological processes. The fastICA [65] algorithm in R was run using the wrapper package picaplot v.0.99.7 (https://github.com/jinhyunju/picaplot). The number of components to be estimated was automatically detected by the implementation using a 70% variance cut-off value. The ICA algorithm was run 15 times and only the components that replicated in every run were used.

##### Pathway-level information extractor (PLIER)

PLIER [9] is a matrix decomposition method that uses prior biological knowledge of pathways and gene sets to deconvolve gene expression profiles as a product of a small number of latent variables (factors) and their gene weights. We performed PLIER analysis using the dedicated R package (v.0.99.0; downloaded from https://github.com/wgmao/PLIER) with the collection of 5,933 gene sets available in the package comprising canonical, immune and chemgen pathways from MSigDB [66], and various cell-type markers from multiple sources. PLIER was run with 100 iterations. Only the 16,440 genes appearing in both gene expression data and the pathway annotation matrix were used. The initial number of latent variables was set using the `num.pc` function provided by the package.

##### Probabilistic estimation of expression residuals (PEER)

PEER [13,67] is a factor analysis method that uses Bayesian approaches to infer hidden factors from gene expression data that explain a large proportion of expression variability. We applied PEER method for co-expression analysis using the peer R package (v.1.0; downloaded from https://github.com/PMBio/peer) with default parameters, accounting also for the mean expression. The initial number of factors was determined using the num.pc function from the PLIER package.

### Functional enrichment analysis

We used the g:GOSt tool from the g:Profiler toolset [29] via dedicated R package gprofiler2 (v.0.1.8) for functional enrichment analysis of gene modules. The short links to the full enrichment results were automatically generated using the parameter `as_short_link = T` in the function `gost`. The results shown in this paper were obtained with data version e99_eg46_p14_55317af.

### *Cis*-eQTL analysis and fine mapping

We performed *cis*-eQTL analysis using the qtlmap (https://github.com/eQTL-Catalogue/qtlmap) Nextflow [68] workflow developed for the eQTL Catalogue project [69]. Briefly, we performed *cis*-eQTL analysis in a +/- 1 Mb window centered around each gene. We used the first six principal components of both the gene expression and genotype data as covariates in the analysis. The eQTL analysis was performed using QTLtools [70].

For *cis*-eQTL fine mapping, we used the Sum of Single Effects (SuSiE) model [27] implemented in the susieR v0.9.0 R package. We performed fine mapping on a +/- 1 Mb *cis* window centered around the lead eQTL variant of each gene. We performed fine mapping on individual-level genotype and gene expression data. Prior to fine mapping, we regressed out six principal components of the gene expression and genotype data from the gene expression data. To identify significant eQTLs for QTL mapping, we performed Bonferroni correction for each gene to account for the number of variants tested per gene and then used Benjamini-Hochberg FDR correction to identify genes with FDR < 0.1. The fine mapping Nextflow workflow for *cis*-eQTLs is available from GitHub (https://github.com/kauralasoo/susie-workflow).

### Gene module *trans*-eQTL analysis and fine mapping

The MatrixEQTL [26] R package (v2.2) was used for *trans*-eQTL analysis to fit a linear model adjusted for sex, batch (where available) and the first three principal components of the genotype data. Before the analysis, the module eigengene profiles were transformed using the inverse normal transformation to reduce the impact of outlier eigengene values produced by some clustering methods. A total of 6,861,056 autosomal genetic variants with minor allele frequency (MAF) > 0.05 were tested. Due to the partial sharing of individuals between cell types and conditions, the eQTL analysis was performed in each cell type and condition separately. To achieve this, the eigenvectors from the integrated approach were split into cell type specific sub-eigenvectors before the analysis. The results from every analytical setting (data partitioning approach (n = 2), co-expression method (n = 5), cell type (n = 9), 90 *trans*-eQTL analyses in total), were then individually filtered to keep nominally significant variant-module associations (P-value < 5×10^-8^).

Next, we applied SuSiE [27] to fine map the nominally significant associations to independent credible sets of variants. For every gene module, we started fine mapping from the lead variant (variant with the smallest association P-value for this module) and used a +/- 500,000 bp window around the variant to detect the credible sets. We continued fine mapping iteratively with the next best nominally significant variant outside the previous window to account for LD and continued this process until no variants remained for the gene module. This procedure resulted in a total of 864 credible sets across all cell types, co-expression analysis methods and data partitioning approaches (integrated and separate).

To aggregate and summarise overlapping associations, we combined all credible sets into an undirected graph where every node represents a credible set of a module from a triplet (data partitioning approach, co-expression method, cell type) and we defined an edge between two nodes if the corresponding credible sets shared at least one overlapping variant (Additional file 1: Figure S2). The graph was constructed using the igraph R package. After obtaining the graph, we searched for connected components, i.e. subgraphs where every credible set is connected by a path, to combine the vast number of results into a list of non-overlapping loci (n = 601). For every component we defined the lead variant by choosing the intersecting variant with the largest average posterior inclusion probability (PIP) value across all the credible sets in the component.

Genes in physical proximity often have correlated expressions levels and could thus manifest as co-expression modules in our analysis. Consequently, if one or more genes in such modules have *cis*-eQTLs, then these *cis* variant-module associations would also be detected by our approach. To differentiate *cis*-acting co-expressions module eQTLs from true *trans* associations, we decided to add an additional filtering step based on gene-level analysis. We performed gene-level eQTL analysis for individual gene expression traits of the 18,383 protein-coding genes and the 601 lead variants. The gene-level eQTL analysis was performed using the MatrixEQTL R package with the same settings and data transformations as in the module-level analysis described above. From every credible set component, we excluded the variant-module pairs together with corresponding credible sets where no *trans* associations (variant-level Benjamini-Hochberg FDR 5%) were included in the module. As *trans*-eQTLs we consider variants that act on distant genes (> 5 Mb away from the lead variant) and genes residing on different chromosomes. After this filtering step we repeated the process of aggregating credible sets and 303 *non-overlapping loci* remained (Additional file 1: Figure S10; Additional file 2).

To further account for the number of co-expression modules tested, we applied both Benjamini-Hochberg false discovery rate (FDR) and Bonferroni correction at the level of each analytical setting (Additional file 1: Figure S10). We applied the FDR 10% threshold to every module - lead variant pair from each of the 90 analytical settings (data partitioning approach, co-expression analysis method, cell type) and if a pair did not pass the threshold we excluded it together with the corresponding credible set(s) from the results. Bonferroni correction was applied in a similar manner with a threshold P-value 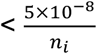, where *n*_*i*_,*i*, = 1,…,90, stands for the number of modules from the corresponding co-expression method and data partitioning approach. We repeated the graph-based aggregation process on the remaining credible sets individually from both correction methods and as a result the FDR 10% threshold reduced the number of significant associations to 140 and Bonferroni threshold to only 4 significant *trans*-eQTLs.

### Colocalisation

We downloaded GWAS summary statistics for 36 blood cell traits [28] from the NHGRI-EBI GWAS Catalog [36]. We downloaded coloc [71] R package v3.1 from bioconda [72]. The *cis*-eQTL colocalisation Nextflow workflow is available from GitHub (https://github.com/kauralasoo/colocWrapper). The same workflow was adjusted for *trans*-eQTL colocalisation.

### Replication of genetic associations

Kim-Hellmuth et al, 2017 [31] profiled gene expression in monocytes before and after stimulation with LPS, muramyl-dipeptide (MDP) and 5′-triphosphate RNA for 90 minutes and 6 hours. We downloaded the eQTL summary statistics from ArrayExpress [53] (accession E-MTAB-5631). Individual-level genotype data were not available for this study. The eQTLGen Consortium [3] performed *trans*-eQTL analysis for 10,317 trait-associated genetic variants in 31,684 whole blood samples. We downloaded the summary statistics from https://www.eqtlgen.org/trans-eqtls.html.

## Declarations

### Ethics approval and consent to participate

Gene expression and genotype data from the CEDAR study were available for download without restrictions from ArrayExpress. For the Fairax_2012, Fairfax_2014 and Naranbhai_2015 studies we applied for access via the relevant Data Access Committee. For the Kasela_2017, we obtained approval from the Data Access Committee of the Estonian Biobank. Ethical approval for the project was obtained from the Research Ethics Committee of the University of Tartu (approval 287/T-14).

### Consent for publication

Not applicable.

### Availability of data and materials

The gene expression matrix, detected gene modules, eigenvectors and *trans*-eQTL credible sets are available in Zenodo (https://doi.org/10.5281/zenodo.3759693). The analysis source code has been deposited to the GitHub repository https://github.com/liiskolb/coexpression-transEQTL.

### Authors’ contributions

LK and KA designed the study. LK performed gene expression data quality control and normalisation, co-expression analysis and *trans*-eQTL analysis. KA performed genotype data quality control and imputation and cis-eQTL analysis. KA and NK wrote Nextflow workflows for the *cis*-eQTL analysis. KA and HP supervised the research. LK and KA wrote the manuscript with input from all co-authors.

## Acknowledgements

We would like to thank Urmo Võsa, Silva Kasela, Kaido Lepik and Sina Rüeger for helpful comments on the manuscript. The computational analyses were performed at the High Performance Computing Center, University of Tartu.

## Supplementary information

**Additional file 1.** Supplementary Figures S1-S10 and Table S1 (PDF)

**Additional file 2.** Co-expression trans-eQTL analysis results together with links to g:Profiler enrichment analysis (XLSX)

## Additional file 1

**Figure S1.**
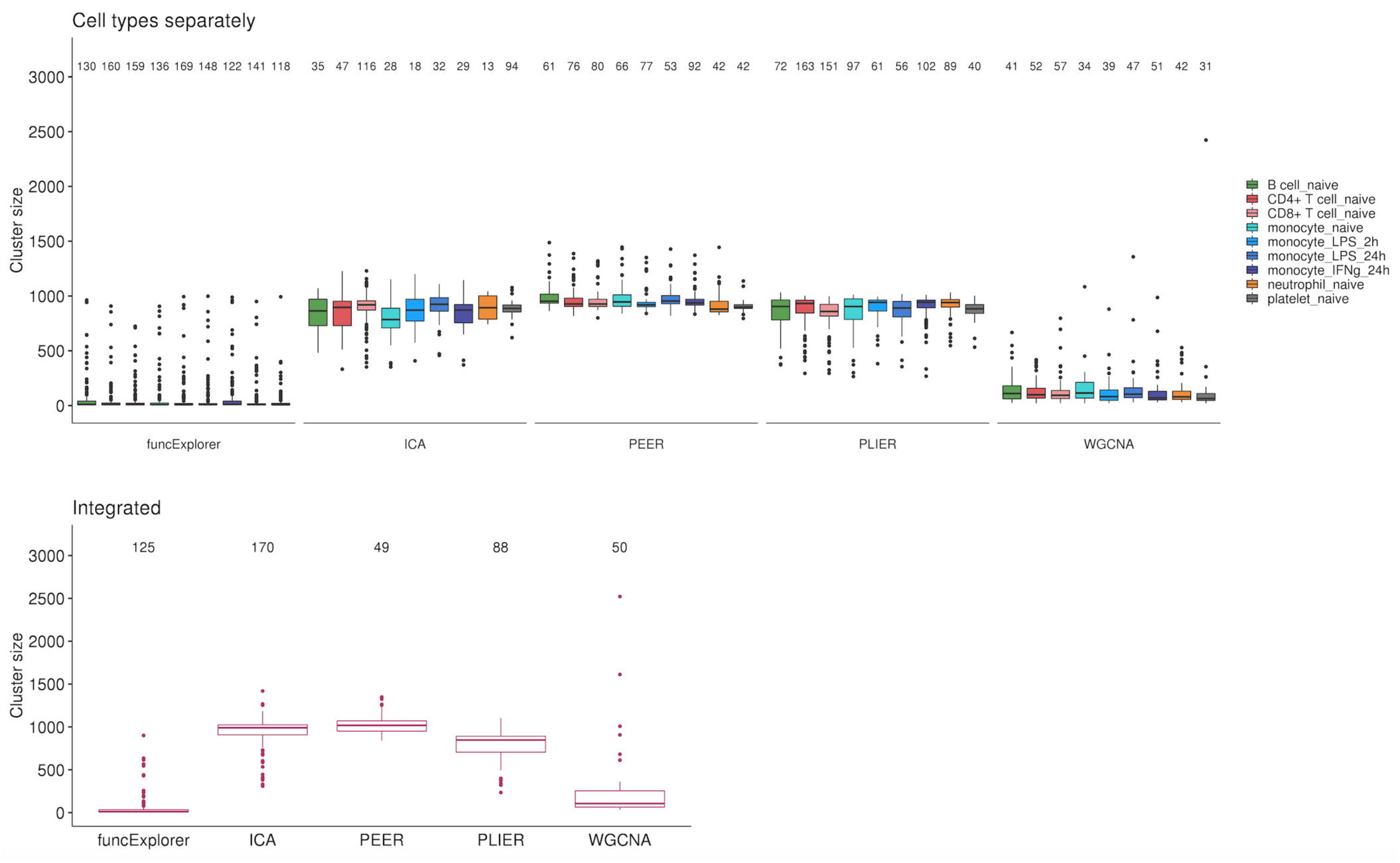
Number of detected gene modules and their size distributions across co-expression analysis methods and data partitioning approaches. The modules from WGCNA and funcExplorer are non overlapping while ICA, PLIER and PEER can define larger modules that share the same set of genes. The numbers on top of the boxplots show the number of modules obtained from each method.

**Figure S2.**
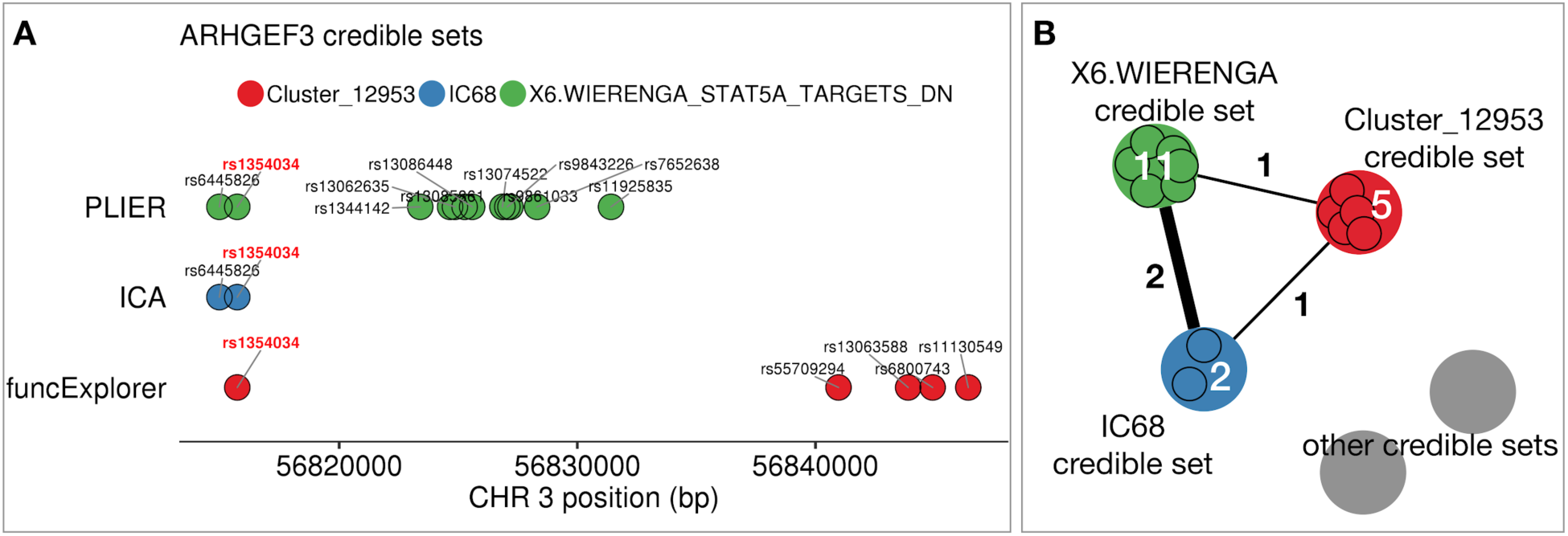
Aggregating fine mapping credible sets in the example of *ARHGEF3 trans*-eQTL locus. We performed fine mapping on the lead variants of every gene module to find corresponding credible sets (see “Methods”). **A** The figure shows an example of three credible sets on chromosome 3 for three gene modules: IC68 from applying ICA to integrated expression data; Cluster_12953 from funcExplorer and X6.WIERENGA_STAT5A_TARGETS_DN from PLIER, both applied to gene expression data from platelets. The associations shown here were detected in platelets. Each circle represents a variant belonging to the corresponding credible set and the different credible sets are distinguished by colors. **B** The credible sets are combined into a graph structure where every node is a credible set of variants and we define an edge between two nodes if they share at least one variant. Grey circles represent all the credible sets from other genomic regions. Here the number of shared variants is shown on the edges and the number of variants belonging to the credible set is shown on the nodes. We searched for connected components, grouped these credible sets together and defined a lead variant for every group based on the largest average posterior inclusion probability (PIP) value within the group. In this example, as all the credible sets share a single variant, rs1354034, then they are grouped together and the overlapping variant with largest PIP is rs1354034.

**Figure S3.**
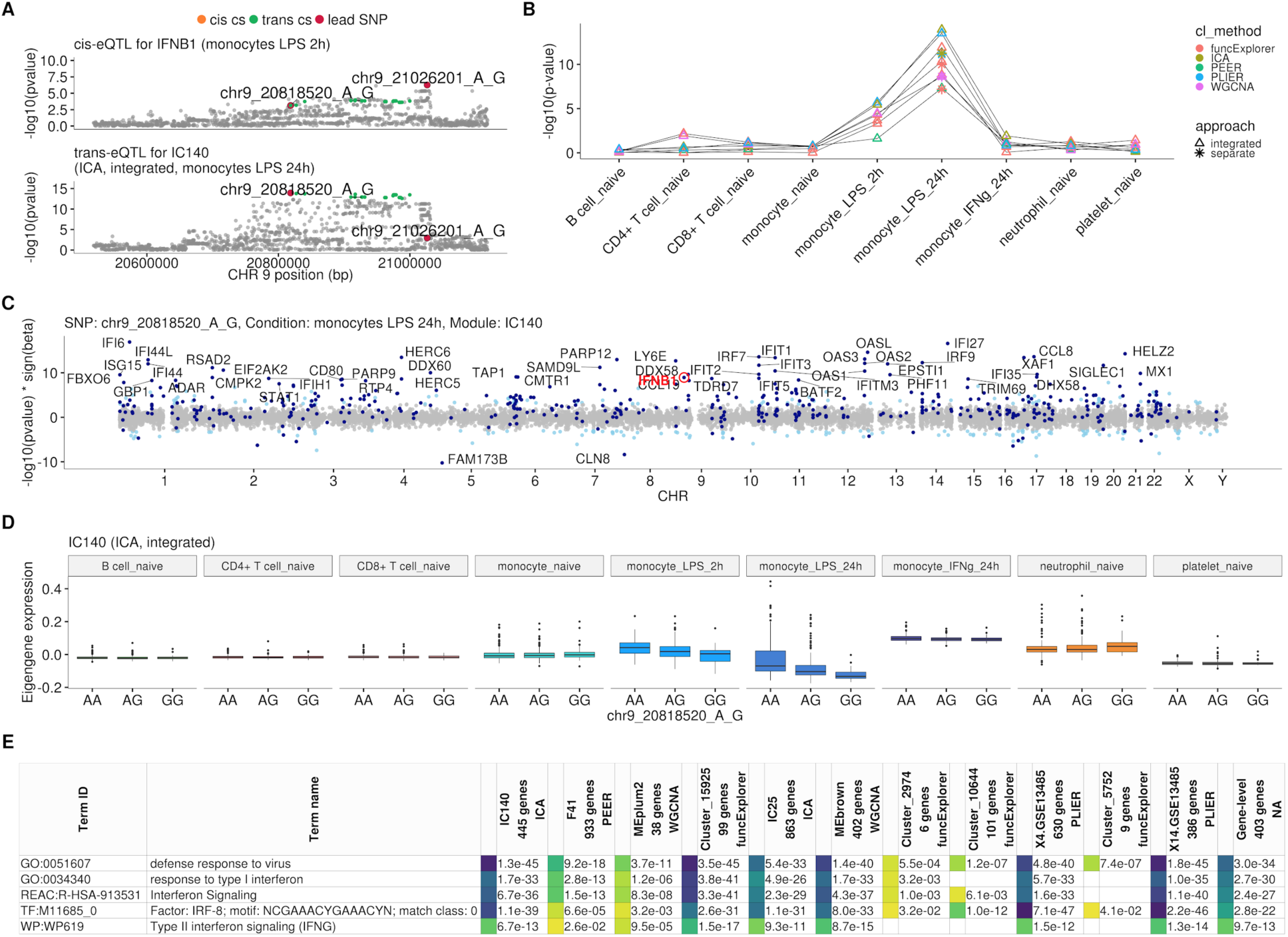
Monocytes specific *trans*-eQTL near *IFNB1* that was first detected by Fairfax et al, 2014 [18] and later replicated in Quach et al, 2016 [17] after 6 hours of LPS stimulation (Additional file 1: Table S1). Both studies performed gene-level *trans*-eQTL analysis only. **A** Association signals for *IFNB1 cis*-eQTL after 2 hours of LPS stimulation and *trans*-eQTL for module IC140 (ICA, 445 genes) after 24 hours of LPS stimulation. We do not detect a colocalisation between the *cis* and *trans* associations, but this could be due to potential confounding caused by probe hybridisation bias in our *cis*-eQTL analysis. **B** Line graph showing that the association between the co-expression modules and *IFNB1 trans*-eQTL lead variant is specific to LPS stimulation. **C** Manhattan plot of gene-level analysis for *trans*-eQTL lead variant rs13296842 (chr9_20818520_A_G). Dark blue points mark genes belonging to module IC140. The module also includes *IFNB1*. **D** Association between *trans*-eQTL lead variant (rs13296842) and eigengene values of a representative ICA module IC140 across cell types. **E** Functional enrichment analysis of the *IFNB1* associated modules (see https://biit.cs.ut.ee/gplink/l/Gi0ZDjn-RM for full results). Empty cell indicates that no gene in the module is annotated to the corresponding term. The last column combines the FDR 5% significant genes from the gene-level analysis. GO - Gene Ontology, REAC - Reactome Pathways, TF - transcription factor binding sites from TRANSFAC, WP - WikiPathways.

**Figure S4.**
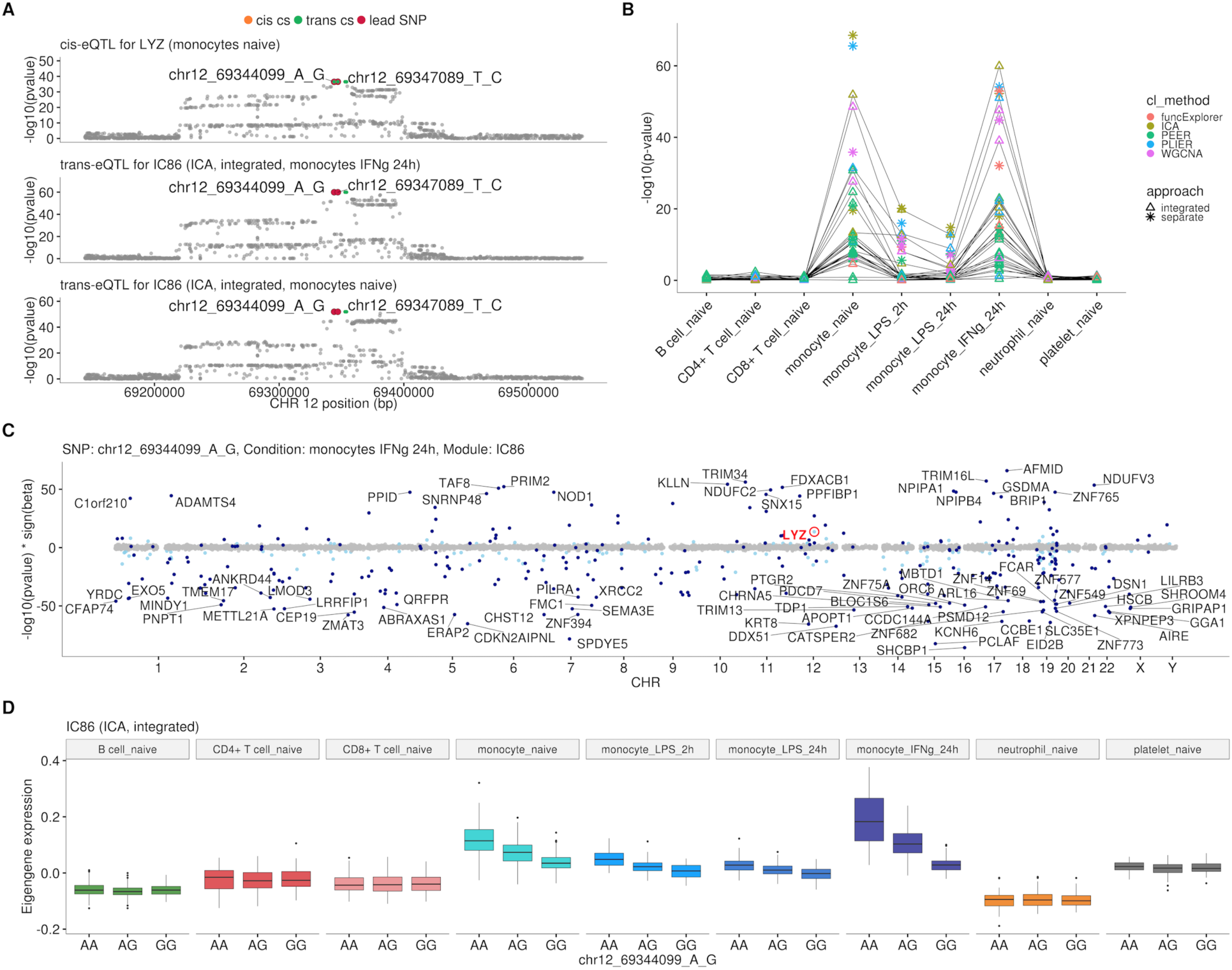
Monocyte-specific *trans*-eQTL near *LYZ* that has been previously detected in three independent studies [10,20,21] (Additional file 1: Table S1). **A** Colocalisation between *cis*-eQTL for *LYZ* in naive monocytes and *trans*-eQTLs for module IC86 (ICA, 309 genes) in naive monocytes and 24 hours after stimulation with IFNγ. The *cis* and *trans* credible sets shown in this figure are identical and contain 4 variants. **B** Line graph showing that the association between the modules and *LYZ trans-*eQTL locus is detected only in naive and stimulated monocytes. This association was detected by at least one module from each of the five co-expression methods and both data partitioning approaches. In total 58 gene modules (54 after Bonferroni threshold) were associated with the locus and 15 of the modules contained the *LYZ* gene itself. There were 14 integrated approach modules that were associated in both naive and stimulated monocytes. **C** Manhattan plot of gene-level analysis for *trans*-eQTL lead variant rs10784774 (chr12_69344099_A_G) in monocytes after 24 h of stimulation with IFNγ. Dark blue points highlight the genes in module IC86. **D** Association between *trans*-eQTL lead variant (rs10784774) and eigengene of ICA module IC86 across cell types. For g:Profiler enrichment results with all associated modules see https://biit.cs.ut.ee/gplink/l/llOr6uCTR8.

**Table S1.** Literature-based replication of *trans*-eQTL loci near *IFNB1, LYZ* and *ARHGEF3* genes. Linkage disequilibrium (r^2^) was calculated using European samples from the 1000 Genomes Phase 3 reference panel.

| trans-eQTL |  |  | Replication |  |  |  |  |  |
| --- | --- | --- | --- | --- | --- | --- | --- | --- |
| Locus | Lead rs ID | Context | Study | Dataset | Context | rs ID | $r^2$ | Replication variant in credible set |
| IFNB1 | rs13296842 | monocytes LPS 24h | Fairfax et al., 2014 [18] | Fairfax_2014 | monocytes LPS 24h | rs2275888 | 0.57 (0.86*) | FALSE |
|  |  |  | Quach et al., 2016 [17] | Quach_2016 | monocytes LPS 6h | rs12553564 | 0.57 (0.86*) | FALSE |
|  |  |  | Ramhdani et al., 2020 [11] | Fairfax_2014 | monocytes LPS 24h | rs2275888 | 0.57 (0.86*) | FALSE |
|  |  |  | Ruffieux et al., 2018 [19] | Fairfax_2014 | monocytes LPS 24h | rs3898946 | 0.88 | TRUE |
| LYZ | rs10784774 | monocytes naive, LPS 2h, LPS 24h, IFNγ 24h | Rotival et al., 2011 [10] | GHS | monocytes | rs11177644 | 0.79 | TRUE |
|  |  |  | Fairfax et al., 2012 [20] | Fairfax_2012 | monocytes | rs10784774 | 1 | TRUE |
|  |  |  | Rakitsch and Stegle, 2016 [21] | CTS | monocytes | rs6581889 | 0.79 | TRUE |
| ARHGEF3 | rs1354034 | platelets | Võsa et al., 2018 [3] | eQTLGen | blood | rs1354034 | 1 | TRUE |
|  |  |  | Mao et al., 2019 [9] | Battle_2014 | blood | rs1354034 | 1 | TRUE |
|  |  |  | Rotival et al., 2011 [10] | GHS | monocytes | rs12485738 | 0.6 | FALSE |
|  |  |  |  |  |  | rs1344142 | 0.6 | TRUE |
|  |  |  | Wheeler et al., 2019 [22] | FHS | blood | - | - | - |
|  |  |  | Nath et al., 2017 [8] | DILGOM07 | blood | rs1354034 | 1 | TRUE |

GHS - Gutenberg Health Study, FHS - Framingham Heart Study, CTS - Cardiogenics Transcriptomic Study, \* - strongest LD in the credible set.

**Figure S5.**
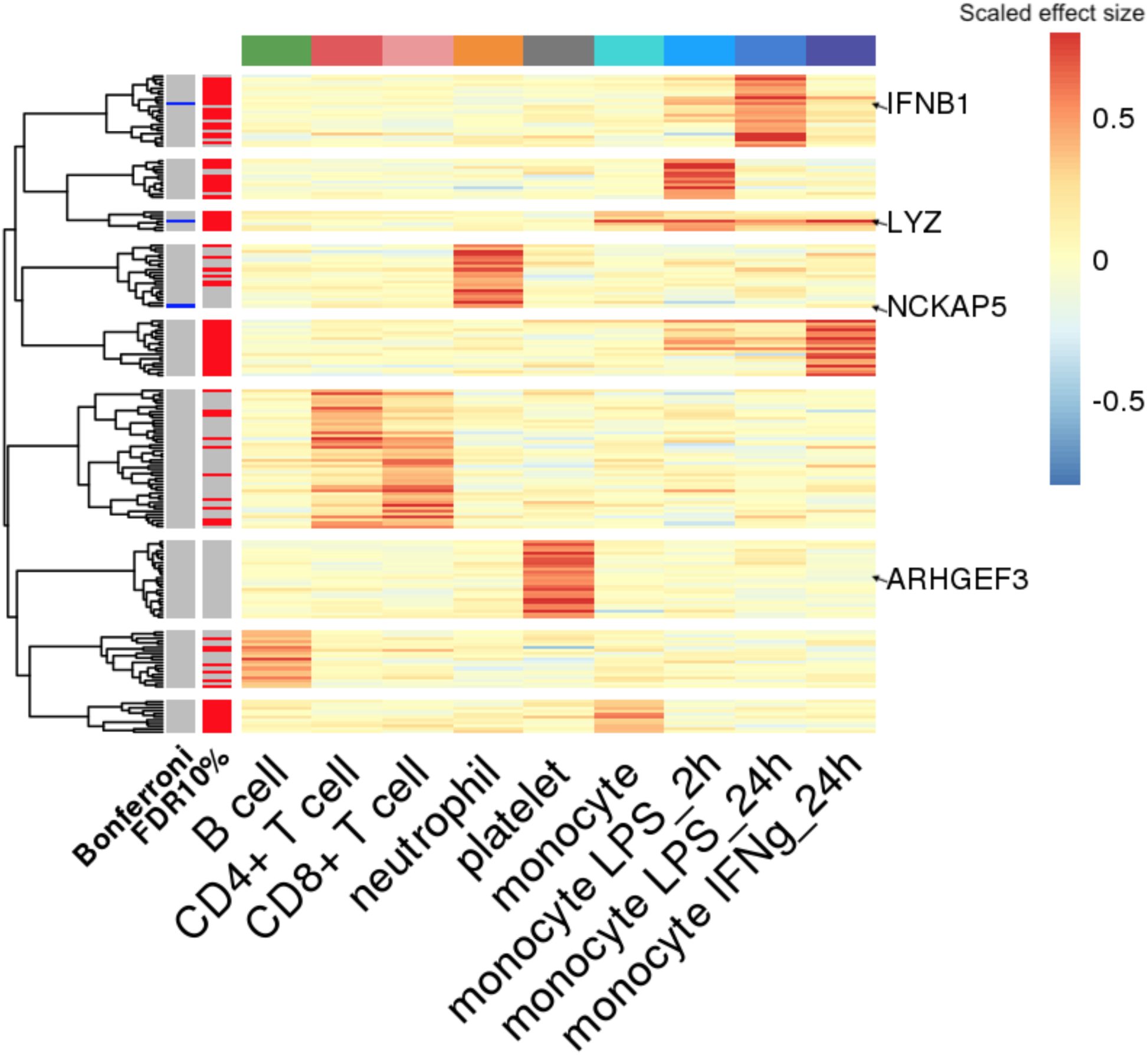
Clustering of *trans*-eQTLs by effect sizes across cell types and conditions. Each locus is represented by the strongest associated gene module from integrated approach, 186 out of the nominally significant 303 loci are shown. Loci associated with modules detected only in cell type and conditions specific data partitions are not shown, because their effect sizes in other cell types are not defined. The effect sizes in every row are scaled relative to the corresponding strongest effect. The annotation bars on the left show indicators for *trans*-eQTLs that were significant after Bonferroni correction (blue stripes) and with FDR 10% (red stripes).

**Figure S6.**
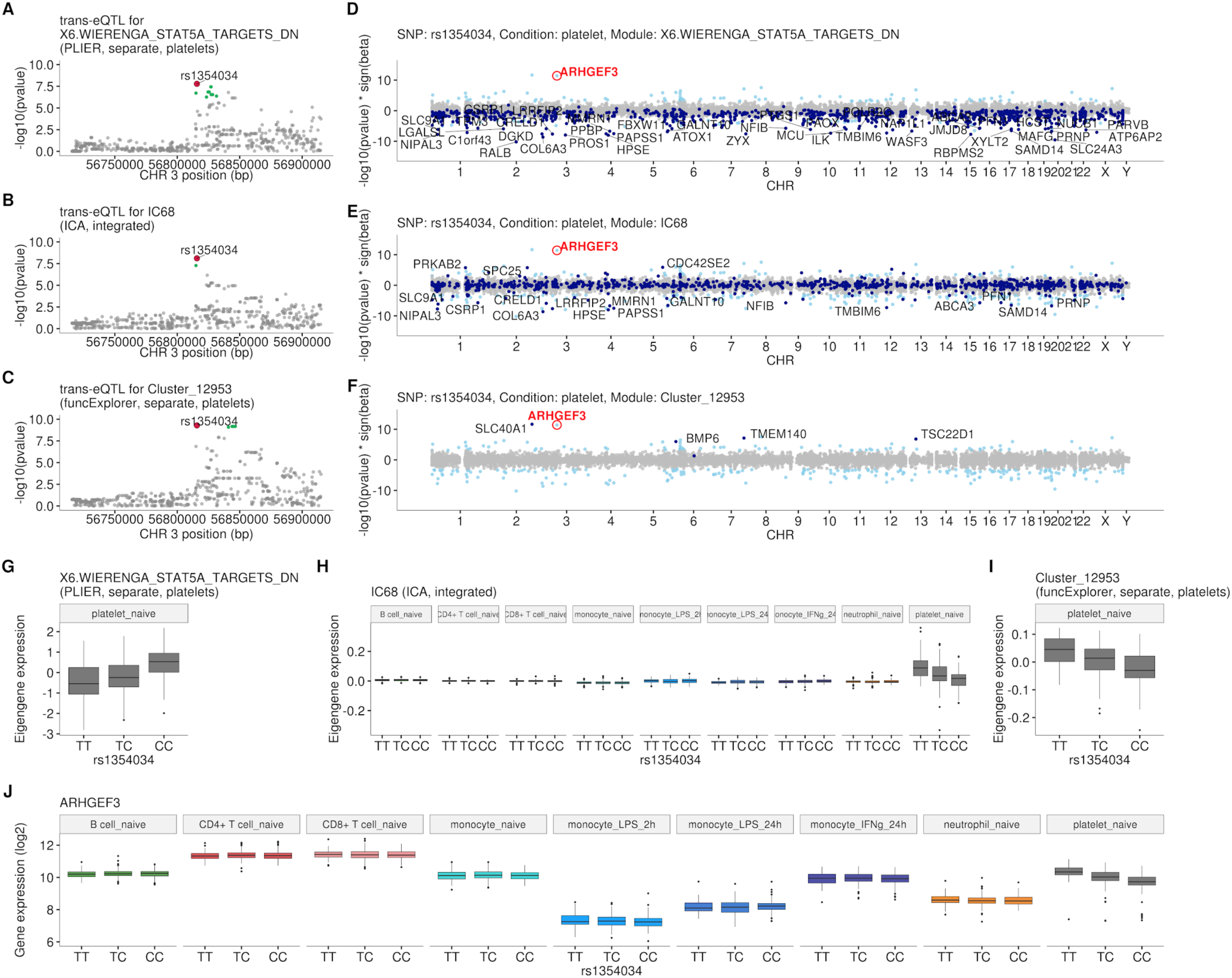
Intronic variant located within the *ARHGEF3* gene (rs1354034) is associated with three co-expression modules in platelets: X6.WIERENGA_STAT5A_TARGETS_DN (918 genes) from PLIER, IC68 (1,074 genes) from ICA and Cluster_12953 (5 genes) from funcExplorer. **A**-**C** Regional plots of *trans*-eQTL summary statistics for each of the modules. Corresponding credible sets are highlighted in green. **D**-**F** Manhattan plots of gene-level eQTL analysis for the *trans*-eQTL lead variant (rs1354034). Dark blue points highlight the genes in the corresponding module. **G**-**I** Association between rs1354034 and eigengene values of three modules. **J** Mean expression of the *ARHGEF3* stratified by the lead variant (rs1354034). Although *ARHGEF3* is expressed in multiple cell types, the *cis*-eQTL effect is only visible in platelets.

**Figure S7.**
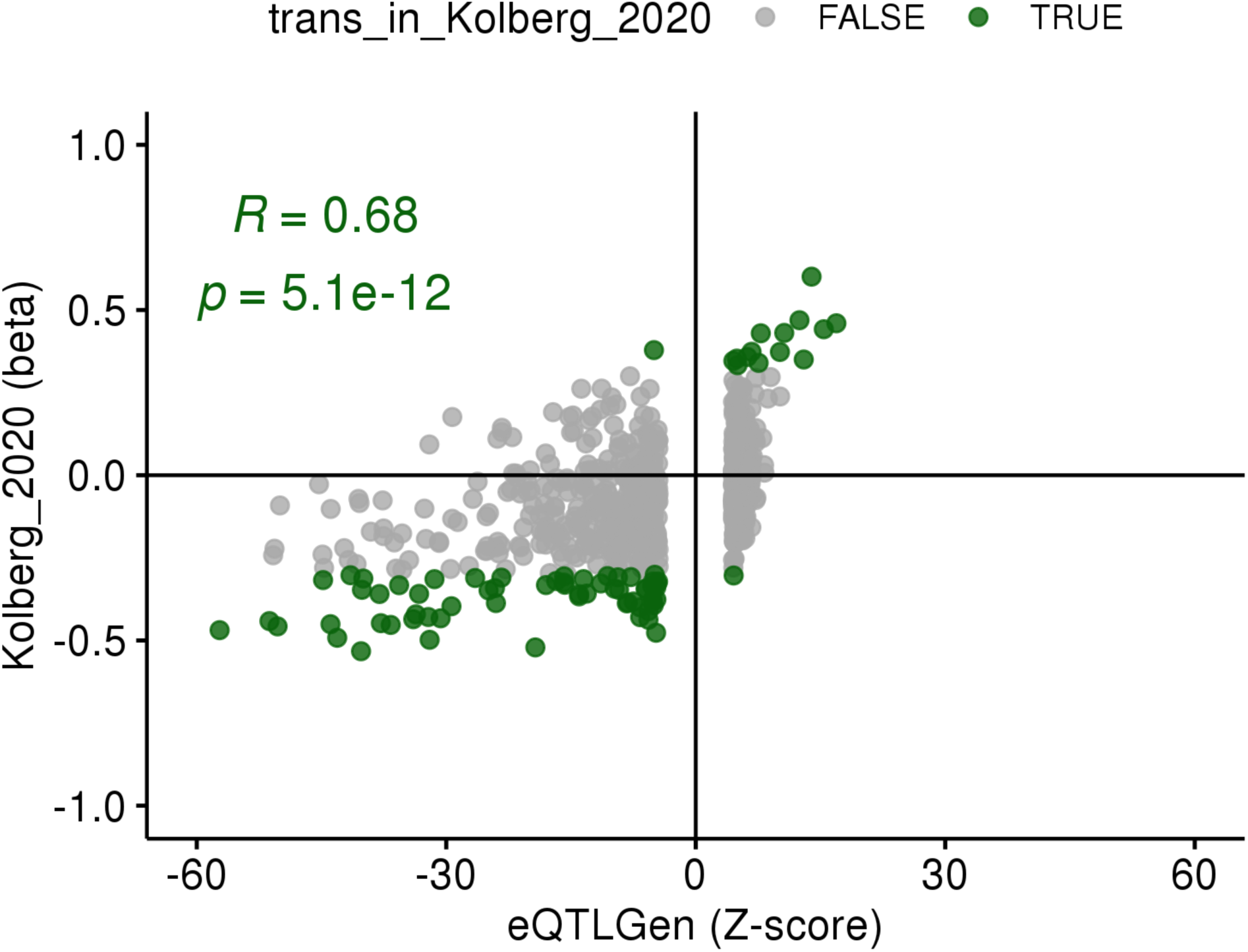
Replication of the *ARHGEF3 trans*-eQTL variant (rs1354034) in eQTLGen [3]. Z-scores of all genes associated with the rs1354034 variant in *trans* from eQTLGen (FDR < 0.05) are shown on the x-axis and effect sizes in platelet samples (n = 226) from this study are shown on the y-axis. The effect sizes for genes significantly associated in platelets (FDR < 0.05, green dots) are correlated between the two studies (Pearson’s r = 0.68) and agree in effect size direction.

**Figure S8.**
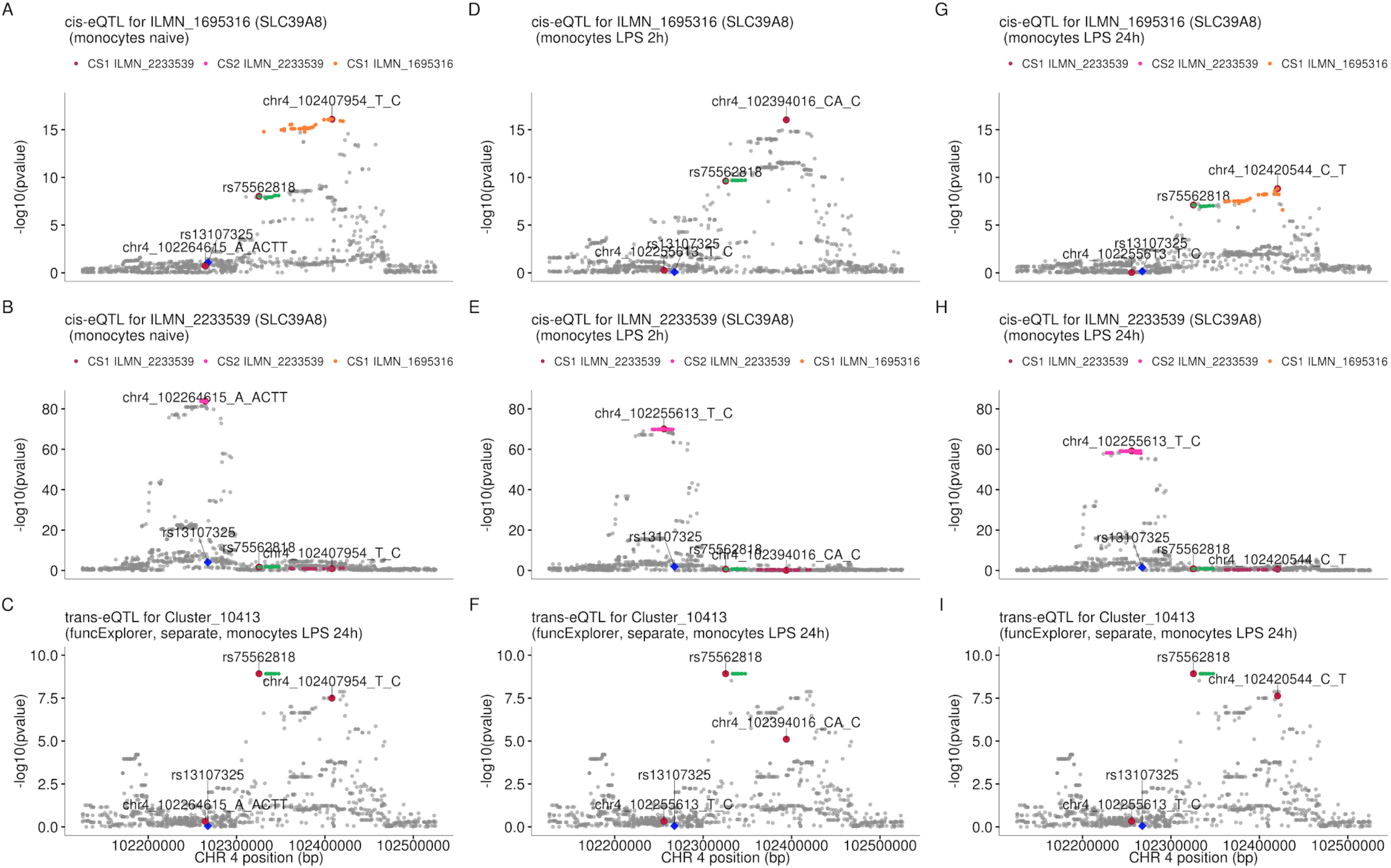
The *SLC39A8 trans*-eQTL lead variant (rs75562818) and *cis*-eQTLs for *SLC39A8* probe sets (ILMN_1695316 and ILMN_2233539) do not colocalise in any of the monocyte conditions in our data. While the *trans* and *cis* credible sets do not overlap, one of the *cis* credible set for probe ILMN_2233539 overlaps with the credible set of probe ILMN_1695316 in naive and 24h LPS stimulated monocytes. **A-C** *cis*-eQTL summary statistics from naive monocytes for probes ILMN_1695316 (**A**), ILMN_2233539 (**B**) and *trans*-eQTL for module Cluster_10413 in monocytes after 24 hours of LPS stimulation (**C**). **D-F** *cis*-eQTL summary statistics from monocytes after 2 hours of LPS stimulation. **G-I** *cis*-eQTL summary statistics from monocytes after 24 hours of LPS stimulation. The *trans* credible set is shown in green. The position of the missense variant in the *SLC39A8* gene (rs13107325) is shown with the blue diamond. The *cis* and *trans* lead variants are highlighted with red circles.

**Figure S9.**
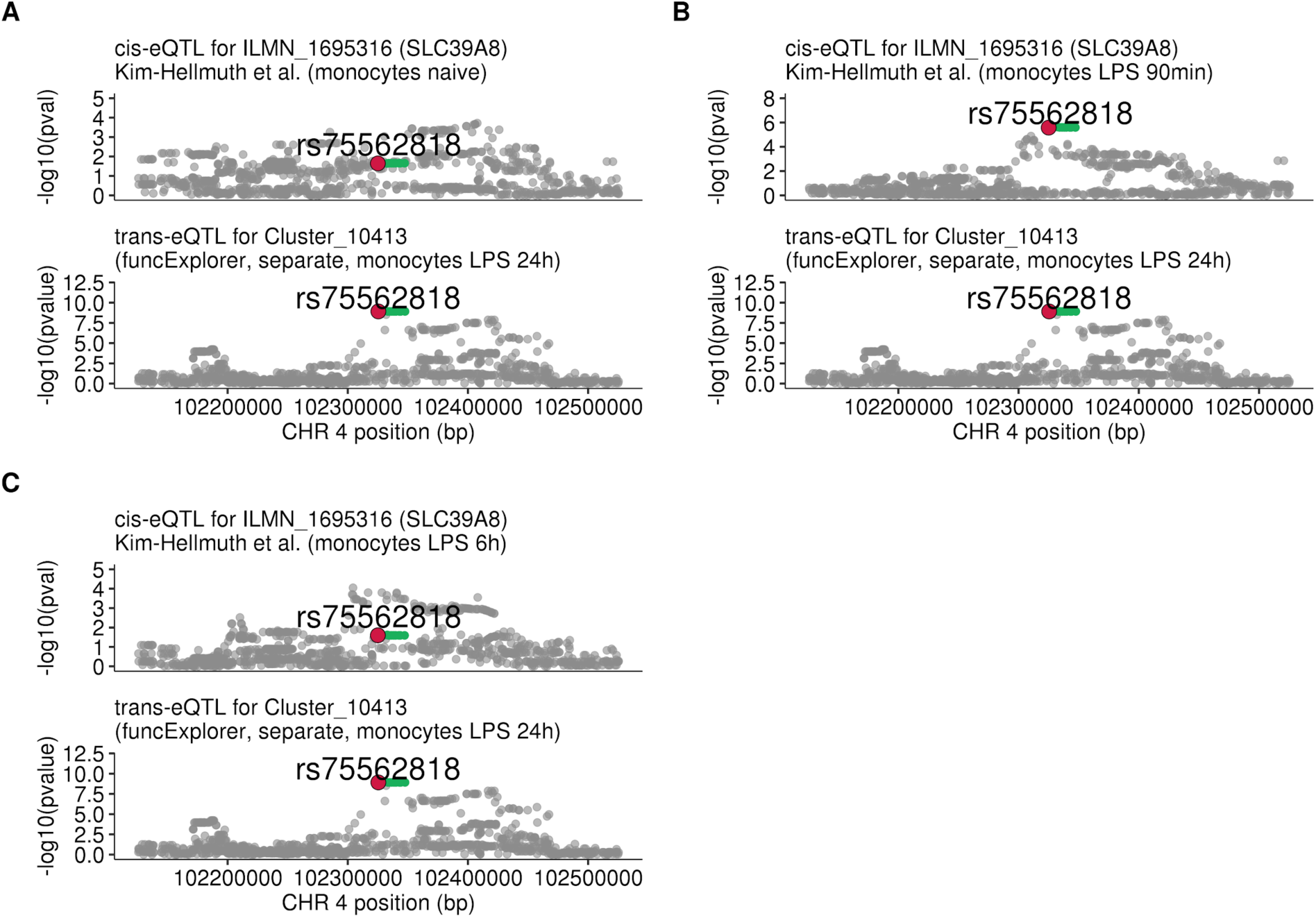
Regional plots comparing association signals of *SLC39A8 trans*-eQTL (rs75562818) and *cis*-eQTLs from Kim-Hellmuth et al, 2017 [31]. **A** *cis*-eQTL summary statistics from naive monocytes (n = 134). **B** *cis*-eQTL summary statistics 90 minutes after LPS stimulation (n = 134). **C** *cis*-eQTL summary statistics 6 hours after LPS stimulation (n = 134). The *cis*-eQTL signal seen after 90 minutes of stimulation has disappeared by 6 hours of stimulation.

**Figure S10.**
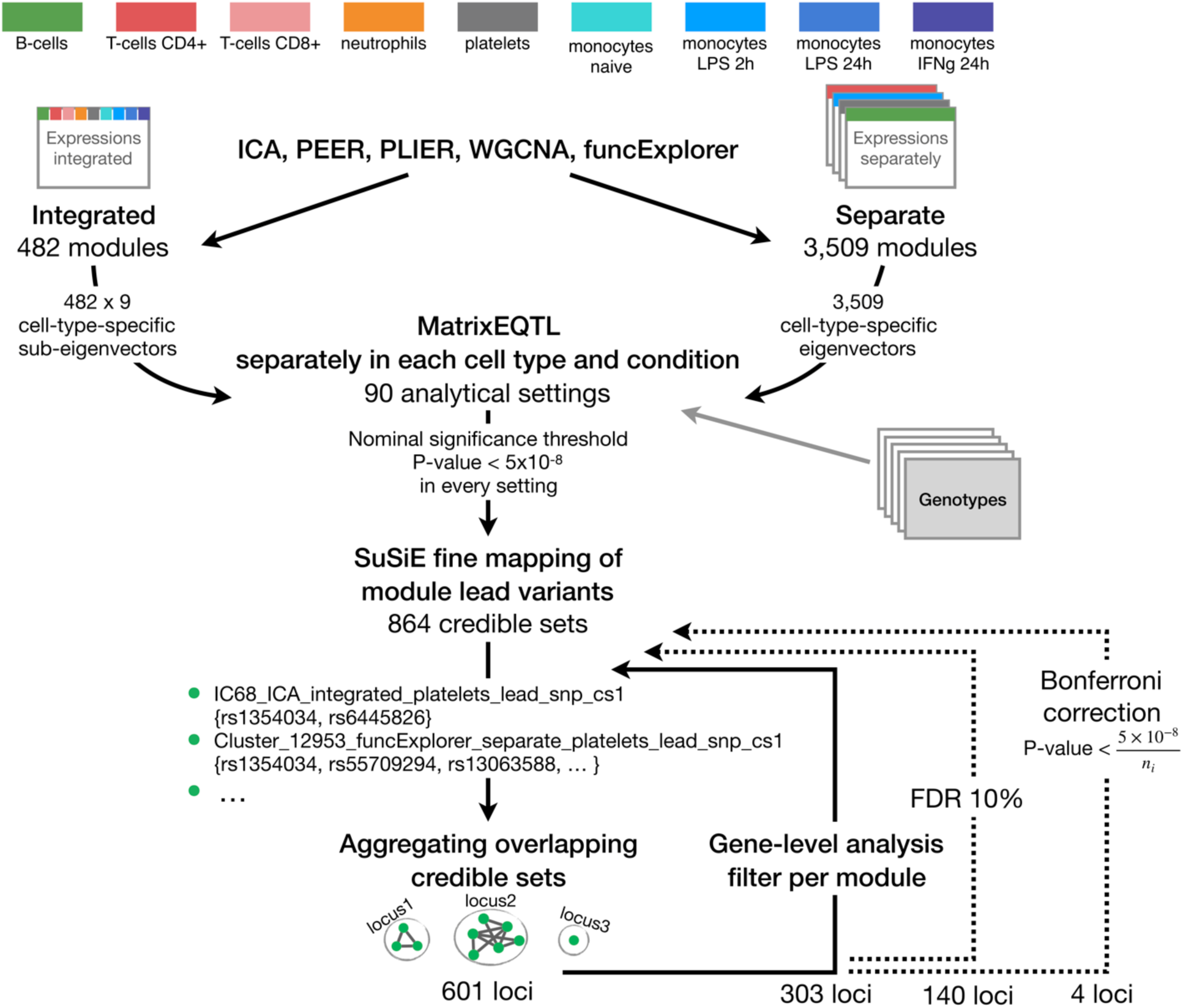
The complete analysis workflow from gene co-expression analysis and gene module *trans*-eQTL mapping to multi-step filtering approaches. Aggregating nominally significant credible sets to non-overlapping groups (Figure S2) resulted in 601 loci. This was followed by gene-level *trans*-eQTL analysis to exclude credible sets of modules where the module does not include any gene-level-significant *trans* genes (FDR 5%). The remaining credible sets were again aggregated using the same approach, 303 loci remained. The credible sets were further filtered by applying FDR 10% threshold to the associations and excluding corresponding credible sets before the aggregation procedure which left us with 140 non-overlapping loci. An even more conservative Bonferroni threshold which takes into account the number of modules detected by corresponding clustering method and analysis setting (n_i_, i = 1,…,90) was applied which further reduced the set of significant *trans*-eQTLs to 4.

